# The assessment of similarity vectors of fingerprint and UMLS in adverse drug reaction prediction

**DOI:** 10.1101/2021.11.15.468509

**Authors:** Milad Beshartifard, Zahra Ghorbanali, Fatemeh Zare-Mirakabad

## Abstract

Identifying and controlling adverse drug reactions is a complex problem in the pharmacological field. Despite the studies done in different laboratory stages, some adverse drug reactions are recognized after being released, such as Rosiglitazone. Due to such experiences, pharmacists are now more interested in using computational methods to predict adverse drug reactions. In computational methods, finding and representing appropriate drug and adverse reaction features are one of the most critical challenges. Here, we assess fingerprint and target as drug features; and phenotype and unified medical language system as adverse reaction features to predict adverse drug reaction. Meanwhile, we show that drug and adverse reaction features represented by similarity vectors can improve adverse drug prediction. In this regard, we propose four frameworks. Two frameworks are based on random forest classification and neural networks as machine learning methods called F_RF and F_NN, respectively. The rest of them improve two state-of-art matrix factorization models, CS and TMF, by considering target as a drug feature and phenotype as an adverse reaction feature. However, machine learning frameworks with fewer drug and adverse reaction features are more accurate than matrix factorization frameworks. In addition, the F_RF framework performs significantly better than F_NN with ACC = %89.15, AUC = %96.14 and AUPRC = %92.9. Next, we contrast F_RF with some well-known models designed based on similarity vectors of drug and adverse reaction features. Unlike other methods, we do not remove rare reactions from the data set in our frameworks. The data and implementation of proposed frameworks are available at http://bioinformatics.aut.ac.ir/ADRP-ML-NMF/.

## 1. Introduction

After the outbreak of coronavirus (SARS-CoV-2), according to studies and performed tests, the World Health Organization (WHO) issued an emergency use authorization for the drug hydroxychloroquine, which was canceled shortly afterward[1]. One of the reasons was the presence of a rare adverse reaction for this drug that caused heart disorders (cardiotoxicity)[2][3]. However, after its widespread use, this drug also caused many deaths[2]. In addition, it has been reported that about 26% of the people are admitted, only in a single hospital in southern India, due to adverse drug reactions (ADRs)[4]. Moreover, drug toxicity is common among children[5] caused to be hospitalized in about 300 children with an average age of 5 years in a medical center in the Netherlands due to ADRs. Such problems show the importance of various drug assessments before a drug is produced and released to the market, considering its time-consuming and costly process.

Nevertheless, sometimes monitoring a drug after launch shows some rare adverse reactions, and it is caused to be withdrawn after a few years, e.g., Rosiglitazone[6]. In May 2007, after examining data from a clinic, it was found that taking Rosiglitazone had a significant effect on the deaths caused by cardiovascular diseases. Eventually, the drug was first withdrawn in Europe, and then in the same year, severe restrictions were imposed on its use in the United States[7].

Regardless of laboratory studies performed at various stages of drug production to identify its adverse reactions, this strategy is still not effective enough to solve the ADR problem. Therefore, there is a severe need to diagnose ADRs accurately. In this regard, researchers are interested in approaching the ADR problem by computational methods. Nowadays, recommender systems[8] and machine learning methods[9][10] have been common computational models for ADR prediction.

A recommender system can predict whether a user prefers an item based on its profile[11]. This technique is also used in the ADR problem. Drugs and adverse reactions are assumed as users and items, respectively. In other words, the recommender system predicts whether a drug has an adverse reaction based on the drug profile[12]. Galeano[13] introduced a collaborative filtering recommendation system to predict an adverse reaction for a new drug using known similar drugs. Matrix factorization[14] is a class of collaborative filtering recommendation systems. Poleksic et al.[15] used the compressed sensing (CS) model as a matrix factorization to predict unknown relationships between drugs and adverse reactions. In addition, the CS model is an appropriate model for dealing with sparsity data. It is suitable for solving the ADR problem because known drug-adverse reactions (positive data) are less than unknowns. In this method, the latent preference of drugs and adverse reactions is computed in a lower dimension by minimizing the defined loss function to complete the drug-adverse reaction associations. Also, later in 2020, Guo et al.[12] recovered drug-adverse reaction matrix using triple matrix factorization (TMF) model based on calculating the similarity between drugs and adverse reactions with different features.

In addition to recommender systems, various machine learning methods are also very effective in solving this problem. Chen et al.[16] predicted the possible likelihood of that drug being associated with adverse effects for each drug. In this method, the similarity of two drugs is calculated based on the interaction of each drug with other drugs and target proteins. Finally, this algorithm uses the other drugs with this adverse reaction to calculate the score of a drug-adverse reaction association based on the relationship between the drugs and calculating the likelihood.

Khan[17] applied different learning models for predicting ADRs such as neural networks, support vector machine, random forest, naive Bayes, using different drug features like fingerprint and drug indications. He limited the SIDER database[18] based on ten adverse reactions with the maximum variance across the drugs. The negative and positive data are defined using the frequency of causing each side effect by a drug according to the recorded medical history of 30 patients in the SIDER database. Therefore, if the frequency of each drug-adverse reaction is more than 0.5, its association is considered positive and vice versa. Finally, for each adverse reaction, it is created a classification model. Zhao et al.[19] proposed a different approach based on the similarity of drugs with different properties such as fingerprints, the two-dimensional structure of drugs, target proteins, ATC code, and some features from the STITCH database. Similarity vectors are considered as the input of the random forest classifier model. Also, the positive data is determined based on known adverse reactions of the drugs with label 1. The negative data is randomly selected from unknown adverse reactions of drugs with label 0. In addition, they chose adverse reactions that are associated with more than five drugs. Rodriguez et al.[20] proposed a Bayesian network approach to predict ADRs using 593 pharmaceutical care center reports. Dey[21] introduced a model to convert two-dimensional or three-dimensional drug structures into numerical vectors using convolutional neural networks (CNNs) for each adverse reaction. Zheng et al.[22] calculated the similarity of drugs based on various properties such as chemical structure, target protein, drug alternatives, and ATC code to form the feature vectors of each drug-adverse reaction association as the model input. Also, they introduced a new approach for selecting negative data. The negative data is chosen based on the assumption that dissimilar drugs have fewer common adverse reactions. Uner[23] proposed a learning method to solve the ADR problem based on CNNs using the structural features of the drug and the characteristics of gene expression. They also selected negative data from pairs of drugs and adverse reactions whose relationship is unknown. In 2020, Liang et al. proposed a new approach for making negative data by random walk on drug-drug interaction networks[24]. They removed adverse reactions associated with less than six drugs from the dataset. In 2021, It was shown that combining different data sources can improve the accuracy of learning models for the ADR problem[25]. Zhang et al.[26] defined negative data based on drug-indication associations and applied a machine learning model to classify adverse reactions as adverse or therapeutic. Table 1 shows a summary of the studies on the ADR prediction problem.

**Table 1:** Some studies on the ADR problem.

| References/Year | Drug Feature | Adverse reaction Feature | Algorithm | Number of Drugs | Number of adverse reactions | Database for adverse reactions | Database for Drug |
| --- | --- | --- | --- | --- | --- | --- | --- |
| 2013[16] | Target protein<br>Literature-Association | - | Machine Learning | 835 | 100 | - | SIDER<br>STITCH |
| 2017[17] | Fingerprint<br>Indication<br>Target Protein | - | Machine Learning | 667 | 10 | - | SIDER<br>ChEMBL |
| 2018[15] | Fingerprint | UMLS semantic | Recommender system | 1430 | 5868 | SIDER<br>MedDRA | SIDER<br>PubChem |
| 2018[19] | Fingerprint<br>Smiles<br>ATC code<br>Target Protein<br>Literature<br>association | - | Machine Learning | 841 | 824 | - | SIDER<br>KEGG<br>RDKit |
| 2018[20] | ADR report | causality categories | Machine Learning | - | 2100 | Northern<br>Pharmacovigilance<br>Centr | - |
| 2019[22] | Fingerprint<br>Target Protein<br>ATC code<br>Substituent | - | Machine Learning | 917 | 500 | - | SIDER<br>DrugBank<br>EMBL-EBI |
| 2019[23] | gene expression<br>SMILES | - | Machine Learning | 791 | 1052 | - | SIDER<br>PubChem |
| 2020[27] | Fingerprint<br>Side effect profile | Drug profile | Recommender System | 614 | 5596 | SIDER<br>PubChem | SIDER |
| 2020[24] | Fingerprint<br>SMILES<br>ATC code<br>Target Protein<br>Literature<br>Association | - | Machine Learning | 841 | 824 | - | SIDER<br>STITCH<br>PubChem<br>KEGG |
| 2021[26] | Indication<br>Target protein | CUI code | Machine Learning | 3632 | 5589 | SIDER<br>-DrugBank | SIDER |

In the computational methods, finding and representing appropriate features of drugs and adverse reactions are challenging. Here, we assess fingerprint and target as drug features; and phenotype and Unified Medical Language System (UMLS) as adverse reaction features to predict adverse drug reaction. Meanwhile, we show that drug and adverse reaction features represented by similarity vectors can improve the performance of the computational methods in adverse drug prediction.

This article proposes four frameworks to analyze drug features and adverse reaction features in drug-adverse reaction association prediction. Two frameworks are based on two machine learning methods; a random forest classifier[10] and a neural network[9]. The rest improve two well-known matrix factorization models; CS [14] and TMF[12].

The first framework is F_RF to predict drug-adverse reactions based on a random forest classifier by considering drug-drug similarity vectors and the vector of adverse reaction similarity. F_RF approach is compared with the neural network-based framework, F_NN, to address the ADR problem. F_RF framework shows better performance than F_NN. Although we improve the CS model[15] into *CS*^*Phen*^ and TMF method[12] into three versions, *TMF*_*Targ*_, *TMF*^*Phen*^ and 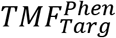 as matrix factorization approaches, the result announces that the F_RF model has better accuracy than state-of-the-art matrix factorization methods. Therefore, we compare the F_RF framework with some well-known algorithms in machine learning approaches that consider similarity vectors as features.

It seems that defining negative data similar to Zheng’s approach[22] and considering similarity vectors as drug and adverse reaction features improve the performance of F_RF. Moreover, unlike some methods[24][19] that remove rare adverse-drug reactions, we consider all drugs and adverse reactions, including rare associations, as training data.

Although the limited association of rare adverse reactions can reduce the model’s accuracy, the F_RF framework achieves comparable performance. Finally, our framework successfully predicts some associations in some case studies.

This paper is organized as follows: the “Materials and methods” section presents the databases and datasets, the notations and definitions, and a description of our proposed frameworks. The “Results” section includes assessing our frameworks and comparing results with other models. The “Discussion” section shows that the F_RF framework successfully predicts adverse reactions for some case study drugs, and finally, the “Conclusion” represents the future point of the ADR problem.

## 2. Material and methods

This section introduces drug and adverse reaction datasets and databases and then defines some notations to describe the problem of the adverse drug reaction (ADR). Finally, the proposed model for ADR prediction is explained in more detail.

### 2.1. Datasets and Databases

In the ADR problem, we need to extract drug and adverse reaction features in addition to known drug-adverse reaction associations from databases. In this paper, we generate three datasets called Δ_1_, Δ_2_, and Δ_3_ from the SIDER database[18], and apply Poleksic[15] and Mizutani[28] datasets as Δ_4_ and Δ_5_, respectively.

We use the SIDER database[18] for collecting adverse reactions (CUI codes), drugs (CID codes), and drug-adverse reaction associations. We split the extracted adverse reactions and drugs from the SIDER database into three datasets, Δ_1_, Δ_2_ and Δ_3_. Fig. 1. shows that the first dataset includes more common adverse reactions in most drugs. In Fig. 2., it can be seen Δ_2_ dataset includes adverse reactions, which are not more common in most drugs. Fig. 3. indicates Δ_3_ dataset contains rare adverse reactions.

**Fig. 1.**
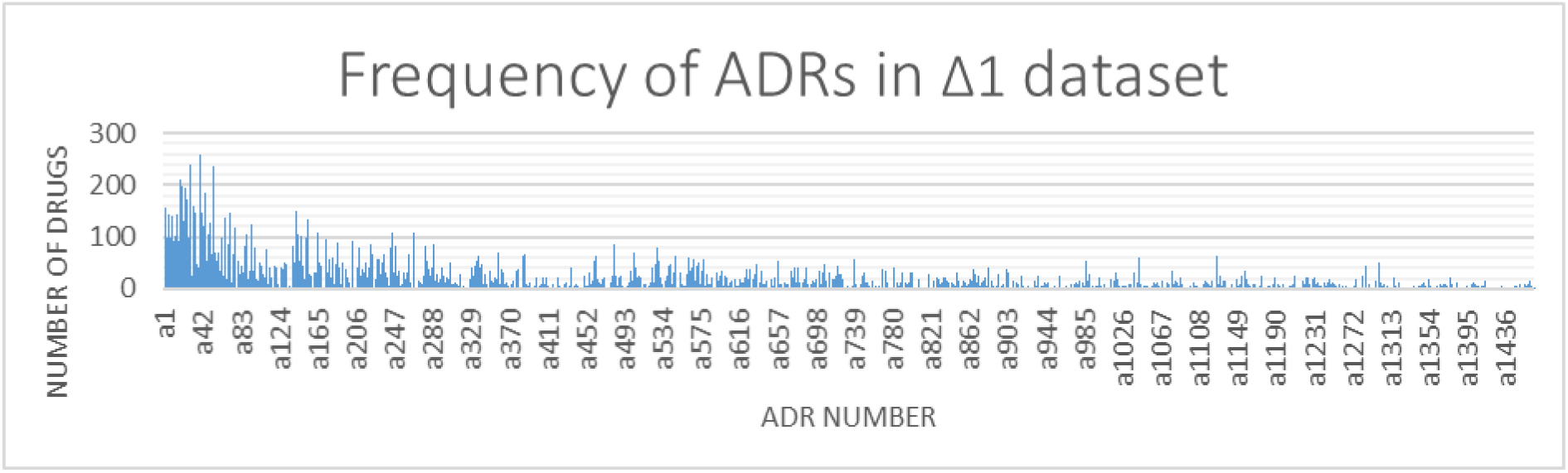
*Frequency of ADRs in* Δ*1 dataset*.

**Fig. 2.**
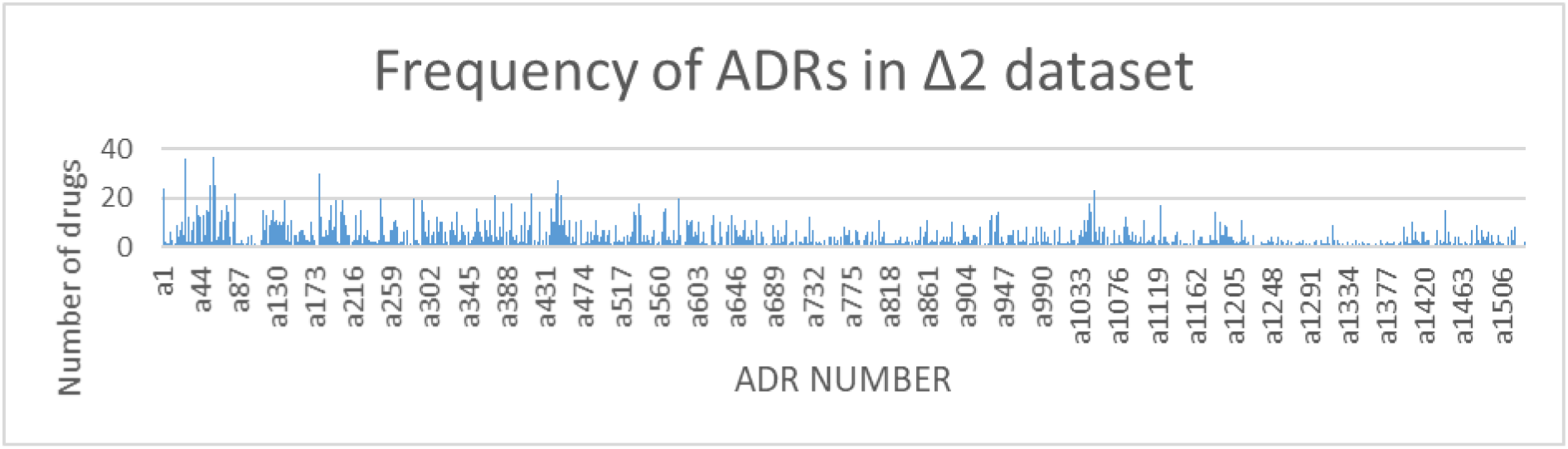
*Frequency of ADRs in* Δ*2 dataset*.

**Fig. 3.**
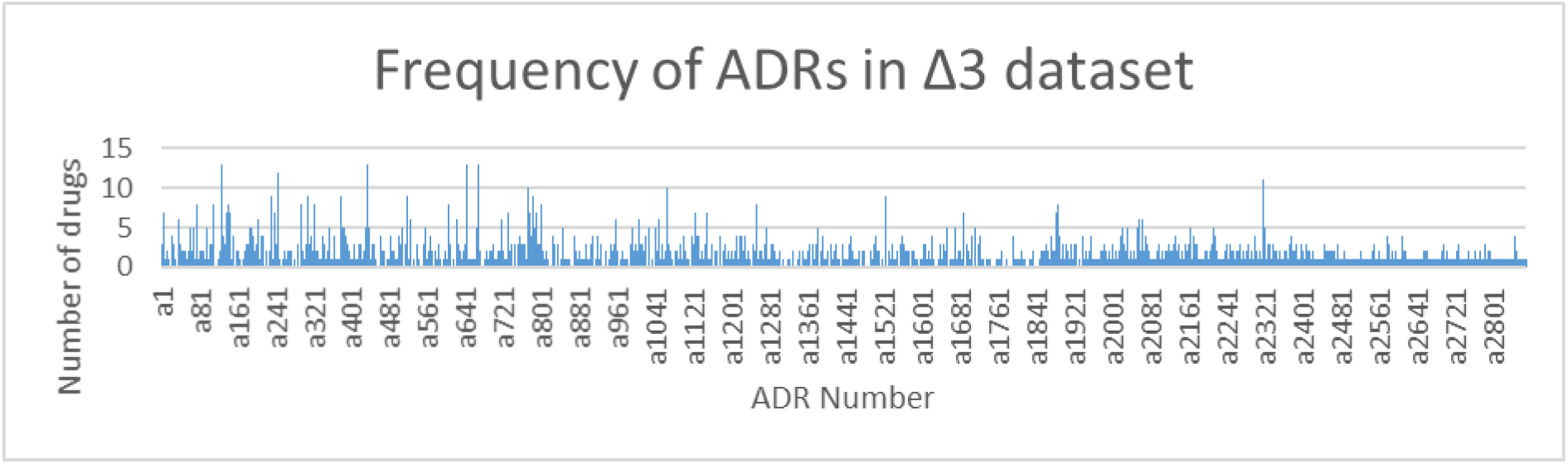
*Frequency of ADRs in* Δ*3 dataset*.

Table 2 shows that dataset Δ ∈ {Δ_1_, Δ_2_, Δ_3_} includes a 2-tuple Δ=< *D*, *A* > where *D* and A indicate the sets of the selected drugs and adverse reactions, respectively. The numbers of drugs and adverse reactions are shown by |*D*| = *m* and |*A*| = *n*, respectively. The first column shows three datasets. The second and third ones display the number of extracted drugs and adverse reactions, respectively, where each adverse reaction has been associated with at least one drug. The fourth column indicates the number of known drug-adverse reaction associations in each dataset. The next column presents the number of adverse reactions that are treated by some extracted drugs. We call treated adverse reactions as indications. Therefore, the set of indications is a subset of *A* as an adverse reaction set. The last column shows the number of drug-indication associations in each dataset.

**Table 2:** Extracted datasets from the SIDER database.

| $\Delta = \langle D, A \rangle$ | $\vert D\vert = m$ | $\vert A\vert = n$ | The number of drug-adverse reaction associations | The number of indications | The number of drug-indication associations |
| --- | --- | --- | --- | --- | --- |
| $\Delta_1$ | 357 | 1467 | 24239 | 663 | 1498 |
| $\Delta_2$ | 443 | 1533 | 4591 | 955 | 3788 |
| $\Delta_3$ | 630 | 2868 | 3017 | 740 | 2786 |

Moreover, we apply Poleksic[15] and Mizutani[28] datasets as Δ_4_ and Δ_5_, respectively. Table 3 shows more details of these datasets. Meanwhile, we extract 17843 and 17418 phenotypes for the adverse reactions in Δ_4_ and Δ_5_ datasets from the CTD database, respectively.

**Table 3:**
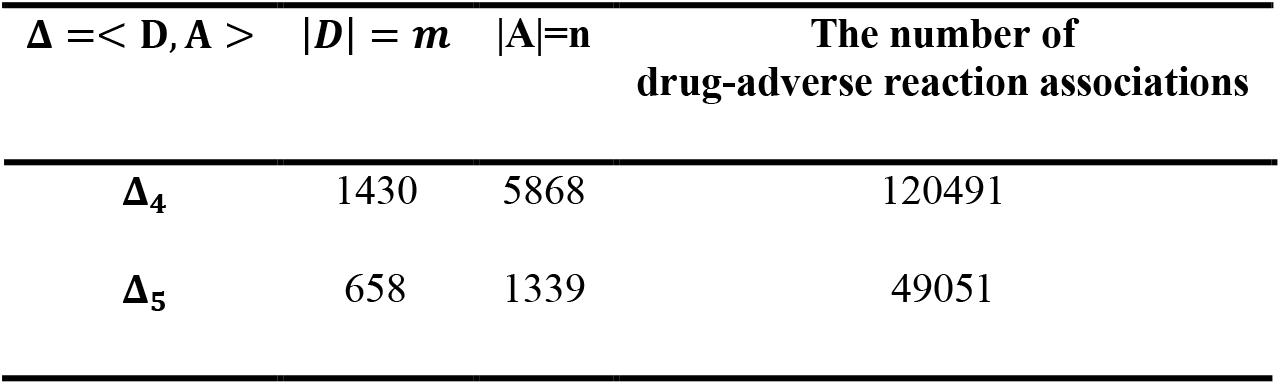
Datasets used in the Poleksic and Mizutani studies.

The fingerprint of each drug as a binary vector with length 881 is extracted from PubChem[25] database. The directly interacted proteins with drugs of the Mizutani database are extracted from DrugBank[29] and Matador[30] databases. The number of these target proteins is equal to 1368.

### 2.2. Notations and definitions

This part describes the selected biological features of a drug and an adverse reaction. Moreover, we define some notations for the feature representations.

#### 2.2.1. Drug

A set of *m* drugs is denoted by *D* = {*d*_1_, *d*_2_, … , *d*_*m*_}, where *d*_*i*_ ∈ *D* shows the *i*^*th*^ drug. Each drug *d* ∈ *D* is displayed by fingerprint chemical structure or protein targets as follows:

- The binary vector *F*^*d*^ = [*f*_1_, … , *f*_881_] with length 881 represents fingerprint[31]. Each *f*_*i*_ with value 1 or 0 represents the existence or absence of the *i*^*th*^ substructure descriptor associated with a specific chemical feature, respectively.
- The binary vector τ^*d*^ = [*τ*_1_, *τ*_2_, … , *τ*_1368_] with length 1368 shows target proteins. Each *τ*_*i*_ with value 1 or 0, represents the *i*^*th*^ protein as a known target for drug *d* or not, respectively.

To calculate the fingerprint chemical structure similarity of drugs *d*, *d*′ ∈ *D*, we use Gaussian Interaction Profile (*GIP*) meter[12][32] defined as follows:

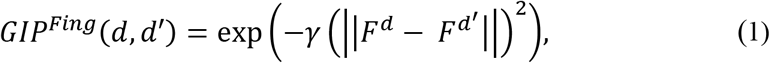

where the bandwidth control parameter (γ) is assigned one[12].

To measure the target protein similarity of drugs *d*, *d*′ ∈ *D*, we apply the cosine similarity (*COS*) criterion[12], as follows:

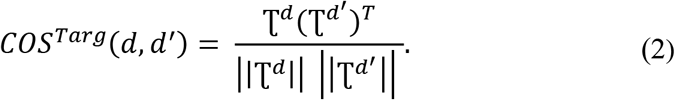

For each drug *d* ∈ *D* and similarity function *f* ∈ {*GIP*^*Fing*^, *COS*^*Targ*^}, we define the similarity vector *δ*^*d,f*^ with length *m*, as follows:

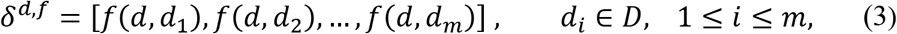

where *δ*^*d,f*^ shows the feature representation of drug *d*.

#### 2.2.2. Adverse reaction

The set of *n* adverse reactions is denoted by *A* = {*a*_1_, *a*_2_, … , *a*_*n*_}, where *a*_*j*_ ∈ *A* shows the *j*^*th*^ adverse reaction. For each *a* ∈ *A*, phenotype or UMLS[33][15] is considered as an adverse reaction feature:

- The phenotype of adverse reaction *a* is shown by the binary vector *P*^*a*^ = [*p*_1_, … . , *p*_18058_]. In our datasets, 18058 is the union of all extracted phenotypes and *p*_*k*_ is considered 1 or 0 to show the existence and absence of the *k*^*th*^ phenotype for adverse reaction *a*. The phenotype illustrates each adverse reaction based on genetic ontology through biological processes, cellular components, and molecular functions[34].
- The UMLS[33] includes over 100 medical terminologies with a unified and semantic network designed by the National Library of Medicine to support scientific research.

To calculate the phenotype similarity of adverse reactions, *a*, *a*′ ∈ *A*, we use the cosine similarity (*COS*) criterion as follows:

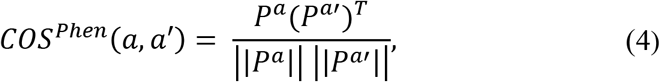

The UMLS similarity of adverse reactions *a*, *a*′ ∈ *A* is computed using UMLS-similarity software which is denoted by function *SIM*^*UMLS*^(*a*, *a*′)[33][15].

For each adverse reaction *a* ∈ *A* and similarity function *f* ∈ {*COS*^*Phen*^, *SIM*^*UMLS*^}, we define the similarity vector α^*a,f*^ with length *n*, as follows:

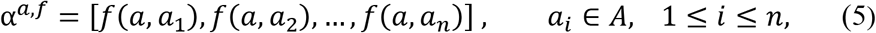

where α^*a,f*^indicates the feature vector of adverse reaction *a*.

### 2.3. Adverse drug reaction problem

We assume that *D* = {*d*_1_, *d*_2_, … , *d*_*m*_} and *A* = {*a*_1_, *a*_2_, … , *a*_*n*_} represent *m* drugs and *n* adverse reactions. In the ADR problem, biological features of drug *d* ∈ *D* and adverse reaction *a* ∈ *A* are given to the model. The primary goal of the ADR problem is to predict the association between adverse reaction *a* ∈ *A* and drug *d* ∈ *D*. If the model predicts adverse reaction *a* associated with drug *d*, the output is one and otherwise zero.

For each drug *d*, we use the similarity vector 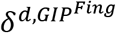, or 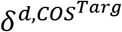 as a drug feature. For each adverse reaction *a*, the similarity vector 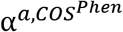 or 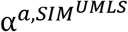 is considered as an adverse reaction feature.

### 2-4. Proposed Models

In this paper, four frameworks are proposed to solve the ADR problem. As machine learning approaches, the first and second frameworks use random forest[10] and neural networks [9], respectively. The third and fourth ones improve CS[15] and TMF[12] as matrix factorization models for ADR prediction. In the following, we describe these frameworks in more detail.

The first and second frameworks are called F_RF and F_NN, respectively. In both models, the concatenation of the similarity vectors 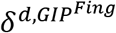 and 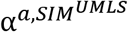 is given as input to predict drug-adverse reaction association. For each drug *d*, the similarity vectors 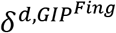 is computed using the GIP function (see Eq.1). Meanwhile, the similarity vector 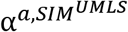 is obtained by the UMLS function[33] for each adverse reaction *a*. In F_RF and F_NN frameworks, we require positive and negative samples for training the model. Known drug-adverse reaction associations and known drug-indication associations are considered positive and negative data, respectively (see Fig. 4.).

**Fig. 4.**
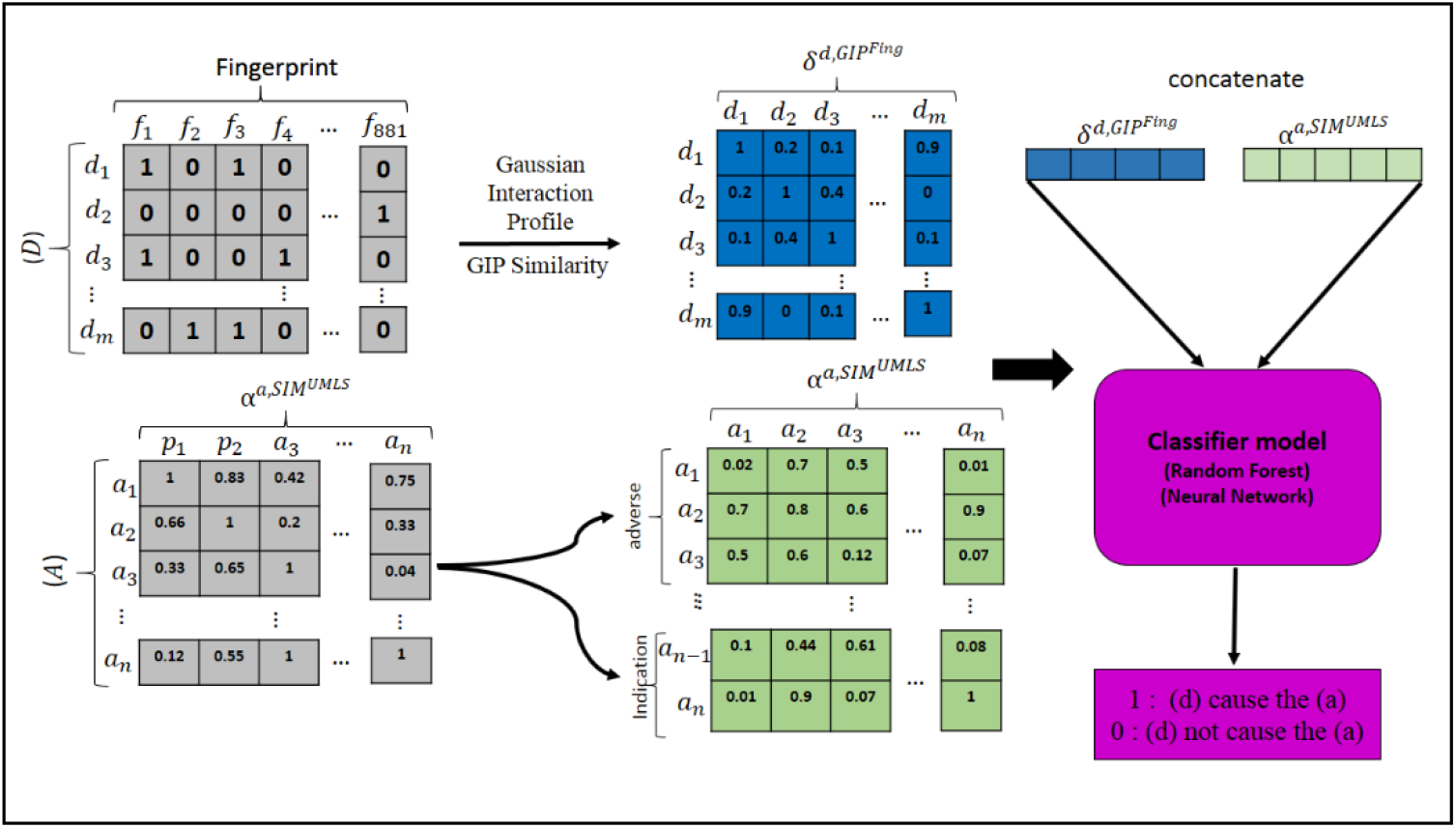
Outline of the first and second frameworks.

The third framework named *CS*^*Phen*^ improves CS[15] model as a matrix factorization approach for ADR prediction. The original CS model uses UMLS-similarity software to calculate adverse reaction similarity for each *a*, *a*′ ∈ *A* shown by *SIM*^*UMLS*^(*a*, *a*′). We improve the similarity matrix between adverse reactions in the CS model by adding the phenotype similarity of adverse reactions, *COS*^*Phen*^ (see Eq.4), to UMLS-similarity as follows:

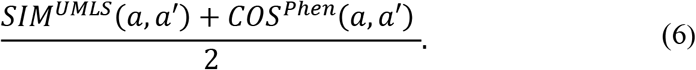

The fourth framework is defined based on TMF[12] model as a matrix factorization approach for ADR prediction. This model combines different criteria to compute the similarity between two drugs *d*, *d*′ ∈. We call this similarity *S*^*TMF*^(*d*, *d*′). Meanwhile, the original TMF uses drug profile as a feature to compute the similarity between two adverse reactions *a*, *a*′ ∈ *A* named *S*^*TMF*^(*a*, *a*′). Here, we define three different versions of TMF as follows:

- *TMF*_*Targ*_: The drug-drug similarity matrix of TMF is changed as:

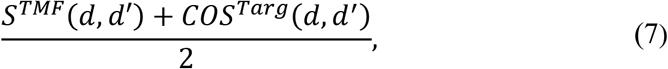

where *COS*^*Targ*^(*d*, *d*′) is obtained based on Eq. 2 to show the similarity between targets of drugs.
- *TMF*^*Phen*^: The similarity matrix between adverse reactions is developed as:

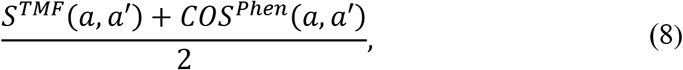

where *COS*^*Phen*^ is obtained based on Eq.4 to show the similarity between phenotypes of adverse reactions.
- 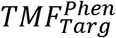: Both drug-drug similarity matrix and the similarity matrix between adverse reactions are improved by Eq.7 and Eq.8.

The matrix factorization models define positive and negative data based on known drug-adverse reaction associations and unknown drug-adverse reaction associations, respectively.

## 3. Result

In this section, we evaluate the four proposed frameworks. The first and second models, F_RF and F_NN, are based on random forest classifier and neural network as machine learning methods. The third framework, *CS*^*Phen*^, improvs CS model as a matrix factorization approach. In the fourth framework, we define three versions of the TMF model, *TMF*_*Targ*_, *TMF*^*Phen*^ and 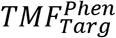, as the matrix factorization model. Each framework was implemented in Matlab 2018b under Windows and Intel Core i5-2430M processor and 4GB of memory.

In the following, we introduce our selected evaluation criteria, then the parameters of each framework are explained. Next, we assess the performance of machine learning frameworks F_RF and F_NN and analyze their effectiveness to predict rare adverse reactions. Then, the assessment of frameworks 3-4, based on matrix factorization, is evaluated. Later, we compare our proposed frameworks to introduce the best one and determine its performance against four well-known models. Finally, we assess our best framework effectiveness on predicting associations of some case studies.

### 3.1. Evaluation criteria

We evaluate our frameworks using the area under receiver operating characteristic curve (AUC), the area under precision-recall curve (AUPRC), and accuracy (ACC) criteria.

AUC is obtained based on the false positive rate (FPR) and the classifier model’s real positive rate (TPR) under different classification thresholds, where:

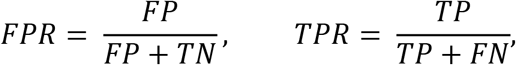

and FP, TN, TP, and FN display false positive, true negative, true positive, and false negative, respectively (see Table 4).

**Table 4:** Definition of true positive (TP), false positive (FP), true negative (TN), and false negative (FN).

| Prediction | Definition |
| --- | --- |
| **True Positive (TP)** | the number of known drug-adverse reaction associations predicted correctly by the model |
| **False Positive (FP)** | the number of drug-indication associations predicted wrongly by the model |
| **True Negative (TN)** | the number of drug-indication associations predicted correctly by the model |
| **False Negative (FN)** | the number of known drug-adverse reaction associations predicted wrongly by the model |

AUPRC shows the relationship between sensitivity (recall) and positive predictive value (precision), where:

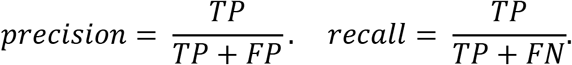

ACC indicates the rate of correct prediction to all predictions as below:

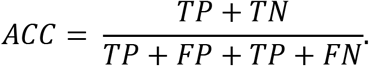

### 3.2. Parameters of the frameworks

The hyperparameters of F_RF, F_NN, *CS*^*Phen*^, *TMF*_*Targ*_, *TMF*^*Phen*^ and 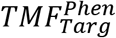 models, respectively, are defined as follows:

- The F_RF framework

In Matlab 2018, the random forest classifier is located in the package Statistics and Machine Learning Toolbox^1^ and has some hyperparameters which can be changed according to the problem. Here, we refer to the three most important ones:

○ “MinLeafSize” shows the minimum observations (samples) per leaf, which is essential in dividing the nodes in the decision trees. By default, this parameter is 1 for classification. A smaller number of “MinLeafSize” makes the model more prone to capturing noise in the training data.
○ “NumPredictorsToSample” means the number of predictor or feature variables to select at random for each decision split. By default, it is equal to the square root of the total number of variables for classification.
○ “NumLearningCycles” variable represents the number of decision trees in the random forest.

As it can be seen in Table 5, we find the best values for these parameters on each dataset Δ∈ {Δ_1_, Δ_2_, Δ_3_} by trial and error.

**Table 5:** Hyperparameters used in the Random Forest model.

| Datasets | Parameters |  |  | Evaluation criteria |  |  |
| --- | --- | --- | --- | --- | --- | --- |
|  | MinLeafSize | NumPredictorsToSample | NumLearningCycles | AUC | AUPRC | ACC |
| $\Delta_1$ | 1 | 6 | 38 | 0.5054 | 0.3201 | 0.7301 |
|  | **1** | **3** | **600** | **0.9637** | **0.9417** | **0.9031** |
|  | **1** | **6** | **38** | **0.9754** | **0.9354** | **0.9288** |
|  | 3 | 6 | 38 | 0.9601 | 0.9440 | 0.9050 |
| $\Delta_2$ | 3 | 6 | 11 | 0.9673 | 0.8292 | 0.9109 |
|  | 1 | 9 | 11 | 0.9540 | 0.8611 | 0.9077 |
|  | 1 | 6 | 42 | 0.9661 | 0.9255 | 0.9132 |
| $\Delta_3$ | **1** | **6** | **38** | **0.9395** | **0.9055** | **0.8799** |

- The F_NN framework

We use a neural network strategy using the back-propagation approach for learning to predict drug-adverse reaction associations. Our model contains an input layer (according to the size of features) followed by one fully connected hidden layer (with100 neurons) and an output layer that decides whether a drug and an adverse reaction are associated or not based on sigmoid function. The learning rate is 0.05, and we train the network about 100 iterations. Our activation function for the hidden layer is tan-sigmoid, and errors are backpropagated according to Scaled Conjugate Gradient (SCG) strategy.

- Improved CS and TMF models

The parameters of these two models are not changed and set according to the original models[12][15].

### 3.3. The assessment of machine learning methods on ADR problem

In this subsection, we assess F_RF and F_NN frameworks on each dataset Δ ∈ {Δ_1_, Δ_2_, Δ_3_}. Moreover, we compare our models to find which one is more accurate on rare adverse reactions.

#### 3.3.1. The assessment of F_RF and F_NN frameworks on Δ_1_, Δ_2_ and Δ_3_ datasets

For each dataset Δ ∈ {Δ_1_, Δ_2_, Δ_3_} (see Table 2), we consider Δ=< *D*, *A* > where D and A indicate drug and adverse reaction sets. All known drug-adverse reaction associations are considered as positive data (*P*^Δ^). Similar to Zhang et al.[26], we suppose that if a drug is prescribed for one of the adverse reactions, it does not cause that adverse reaction. According to this assumption, we extract drug-indication associations as negative data (*N*^Δ^).

To form a test set from the dataset, we randomly select 10% of negative data 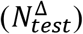 and the exact size of positive data 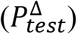. The rest of the positive 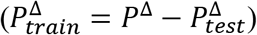 and negative data 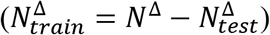 are considered as the training data (see Table 6).

**Table 6:** the number of training and test data in each dataset.

| Datasets | Associations in dataset |  | Associations in training data |  | Associations in test data |  |
| --- | --- | --- | --- | --- | --- | --- |
| | $\vert P^\Delta\vert $ | $\vert N^\Delta\vert $ | $\vert P_{train}^\Delta\vert $ | $\vert N_{train}^\Delta\vert $ | $\vert P_{test}^\Delta\vert $ | $\vert N_{test}^\Delta\vert $ |
| $\Delta_1$ | 24239 | 1498 | 24089 | 1348 | 150 | 150 |
| $\Delta_2$ | 4591 | 3788 | 4212 | 3409 | 379 | 379 |
| $\Delta_3$ | 3017 | 2786 | 2738 | 2507 | 279 | 279 |

As it can be seen in Table 6, the training data extracted from Δ_1_ dataset is imbalanced because the number of negative data is less than positive data. We balance this training data by oversampling strategy to repeat the negative data.

For each dataset Δ, we train F_RF and F_NN frameworks on 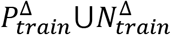 and test on 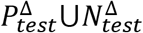. Table 7 shows the values of corresponding evaluation criteria on the test set. Both frameworks have better performance on Δ_1_ dataset because the number of training data in this dataset is more than the others with common adverse reactions. The average accuracy on all evaluation criteria on F_RF is more than in the F_NN framework.

**Table 7:** The evaluation criteria of F_RF and F_NN frameworks on the test set.

| $F\_RF(P_{test}^{\Delta} \cup N_{test}^{\Delta})$ | AUC | AUPRC | ACC | $F\_NN(P_{test}^{\Delta} \cup N_{test}^{\Delta})$ | AUC | AUPRC | ACC |
| --- | --- | --- | --- | --- | --- | --- | --- |
| $\Delta_1$ | **0.9637** | **0.9401** | **0.9032** | $\Delta_1$ | **0.9735** | **0.9486** | **0.9132** |
| $\Delta_2$ | 0.9612 | 0.9076 | 0.8879 | $\Delta_2$ | 0.9344 | 0.9247 | 0.8694 |
| $\Delta_3$ | 0.9592 | 0.9392 | 0.8835 | $\Delta_3$ | 0.9308 | 0.9050 | 0.8513 |
| **Average** | 0.9614 | 0.9290 | 0.8915 | **Average** | 0.9462 | 0.9261 | 0.8780 |

#### 3.3.2. The assessment of F_RF and F_NN frameworks on the rare adverse reactions

An adverse reaction associated with a maximum of two drugs (and a minimum of one drug) is considered as adverse reaction rare. Predicting rare adverse reactions is an obstacle because the known associations of them are too limited. Some studies[24][19] exclude the rare adverse reactions from their dataset to increase their performance. The number of drug–rare adverse reaction associations in each dataset Δ_1_, Δ_2_ and Δ_3_ contain 582, 668, and 1797, respectively. Fig. 1., Fig. 2., and Fig. 3. depict the ratio of the adverse reaction numbers and their related drugs in each dataset Δ ∈ {Δ_1_, Δ_2_, Δ_3_}. Each adverse reaction in datasets Δ_1_, Δ_2_, and Δ_3_ is associated with average 16.52, 2.99, and 1.05 drugs, respectively.

To evaluate the accuracy of F_RF and F_NN frameworks on rare adverse reactions in Δ dataset, we choose positive test data 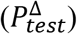 with 10% drug-adverse reaction associations known as rare adverse reactions. The sets of 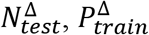 and 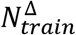 are generated the same as the previous section. Table 8 illustrates the results for predicting rare adverse reactions on the test set. In both models, the performance of Δ_3_ dataset is better than other ones because it has more rare adverse reactions than the other datasets. The results show that F_RF is generally more accurate than the F_NN framework.

**Table 8:** The evaluation of F_RF and F_NN frameworks on rare adverse reactions as the test set.

| $F\_RF(P_{test}^{\Delta} \cup N_{test}^{\Delta})$ | AUC | AUPRC | ACC | $F\_NN(P_{test}^{\Delta} \cup N_{test}^{\Delta})$ | AUC | AUPRC | ACC |
| --- | --- | --- | --- | --- | --- | --- | --- |
| $\Delta_1$ | 0.8390 | 0.7465 | 0.7400 | $\Delta_1$ | 0.8733 | 0.8456 | 0.8212 |
| $\Delta_2$ | 0.9136 | 0.8833 | 0.8298 | $\Delta_2$ | 0.8702 | 0.8534 | 0.7902 |
| $\Delta_3$ | **0.9345** | **0.9182** | **0.8428** | $\Delta_3$ | **0.9101** | **0.9015** | **0.8127** |
| **Average** | 0.8957 | 0.8493 | 0.8042 | **Average** | 0.8845 | 0.8668 | 0.8080 |

### 3.4. The assessment of matrix factorization methods on ADR problem

In this subsection, we assess the performance of the improved CS model[15] called *CS*^*Phen*^, and TMF model[12] in three versions, *TMF*_*Targ*_, *TMF*^*Phen*^ and 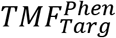.

As mentioned above, these models are extended by adding the new drug and adverse reaction features to the original ones. The main version of these models are performed on datasets Δ_4_[15] and Δ_5_ [28], respectively. Therefore, we perform the improved CS model, *CS*^*Phen*^, and three different versions of the TMF model, *TMF*_*Targ*_, *TMF*^*Phen*^ and 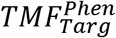, on these datasets.

In these models, positive and negative data are defined based on all known drug-adverse reaction associations and unknown drug-adverse reaction associations, respectively.

To evaluate improved models, we perform cross-validation similar to the original version of CS[15] and TMF[12]. Here, we divide our known drug-adverse reaction associations randomly into equal subsets. One of them is chosen randomly, and its associations are set as 0 in the drug-adverse reaction associations matrix, called the test set. Then, the model is trained by the remaining subsets. For prediction evaluation, the test set is added to the whole matrix as positive samples again. This process is repeated for every subset.

Table 9 depicts the evaluation scores for general CS and TMF models and their improved versions.

**Table 9:** The evaluation of improved and original CS and TMF models.

| Model (*dataset*) | AUC | AUPR |
| --- | --- | --- |
| $CS(\Delta_4)$ | 0.9412 | 0.5059 |
| $CS^{phen}(\Delta_4)$ | **0.9526** | **0.5517** |
| $TMF(\Delta_5)$ | 0.9415 | 0.7071 |
| $TMF^{phen}(\Delta_5)$ | 0.9447 | 0.7093 |
| $TMF_{targ}(\Delta_5)$ | 0.9436 | 0.7100 |
| $TMF_{targ}^{phen}(\Delta_5)$ | **0.9479** | **0.7104** |

### 3.5. Comparison machine learning and matrix factorization methods

This paper makes two similarity matrices of drugs and adverse reactions based on different features of drugs and adverse reactions. Meanwhile, we design four frameworks. Two of them are based on machine learning, F_RF and F_NN, and the others are based on matrix factorization, improvement of CS[15], and TMF[12].

In machine learning frameworks, the contention of each row of similarity matrices is given as input to F_RF and R_NN. In matrix factorization frameworks, the similarity matrices are integrated into the original similarity matrices of CS and TMF approaches.

The results show that although we improve the CS and TMF models as matrix factorization approaches, F_RF and R_NN represent better performance as machine learning approaches. In addition, fewer features are considered for drug and adverse reactions to computing the similarity in machine learning frameworks (see Table 7 and Table 9).

According to corresponding results of applying F_RF and F_NN on datasets (see Table 7 and Table 8), random forest performs more accurately on datasets and predicts rare adverse reactions with higher performance.

### 3.6. Comparison with Related Studies

We compare the best-proposed framework, F_RF, with three machine learning models[24][19][22] that predict drug- adverse reaction associations using similarity-based methods. In addition, we compare F_RF with the logistic regression model introduced by Zhang in 2021[26]. This model defines negative data as ours by considering drug-indication associations for negative data[26]. Table 10 illustrates the values of evaluation criteria for each one.

**Table 10:**
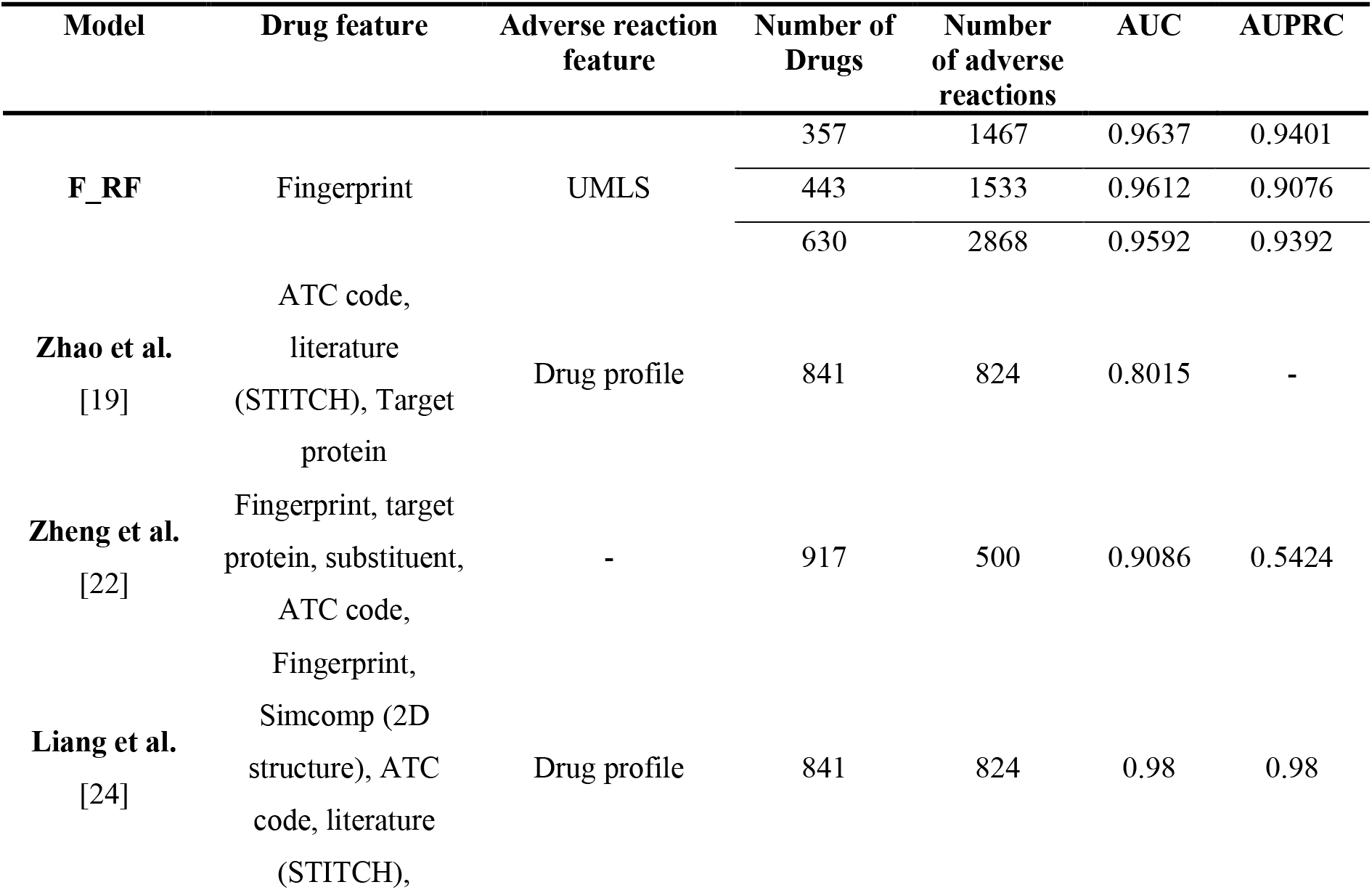
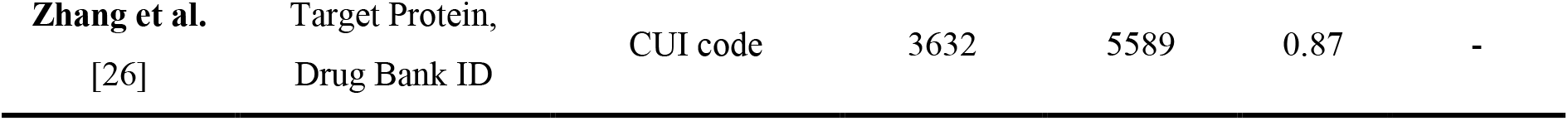
Comparison of the proposed model with other related studies.

It should be noted, the Liang et al. model[24] excluded all adverse reactions with less than six associations from its dataset, and it positively affected the performance. However, the performance of F_RF on datasets includes rare adverse reactions. In addition to considering rare adverse reactions, F_RF uses fewer features than the others while the results are competitive with the rest models.

Moreover, the performance of F_RF is more accurate than Zheng et al.[22], which performs a classifier for each adverse reaction separately.

## 4. Discussion

We also use several antiviral drugs out of our drug set to test the performance of the F_RF model. It should be noted, the association of these drug–adverse reactions are not available in the training set. We assume that positive association is determined based on the predicted probability of an association between a drug-adverse reaction which is more than 0.5. In addition, we check these pairs with F_NN, too.

For drug darunavir, F_RF predicts two adverse reactions Cardiac arrest, a type of heart disorder, and Hepatic steatosis, a type of liver disorder with the probability of association 0.98 and 0.61 (see Table 11). it means darunavir causes these adverse reactions. According to [35][36], the model correctly makes the prediction.

**Table 11:**
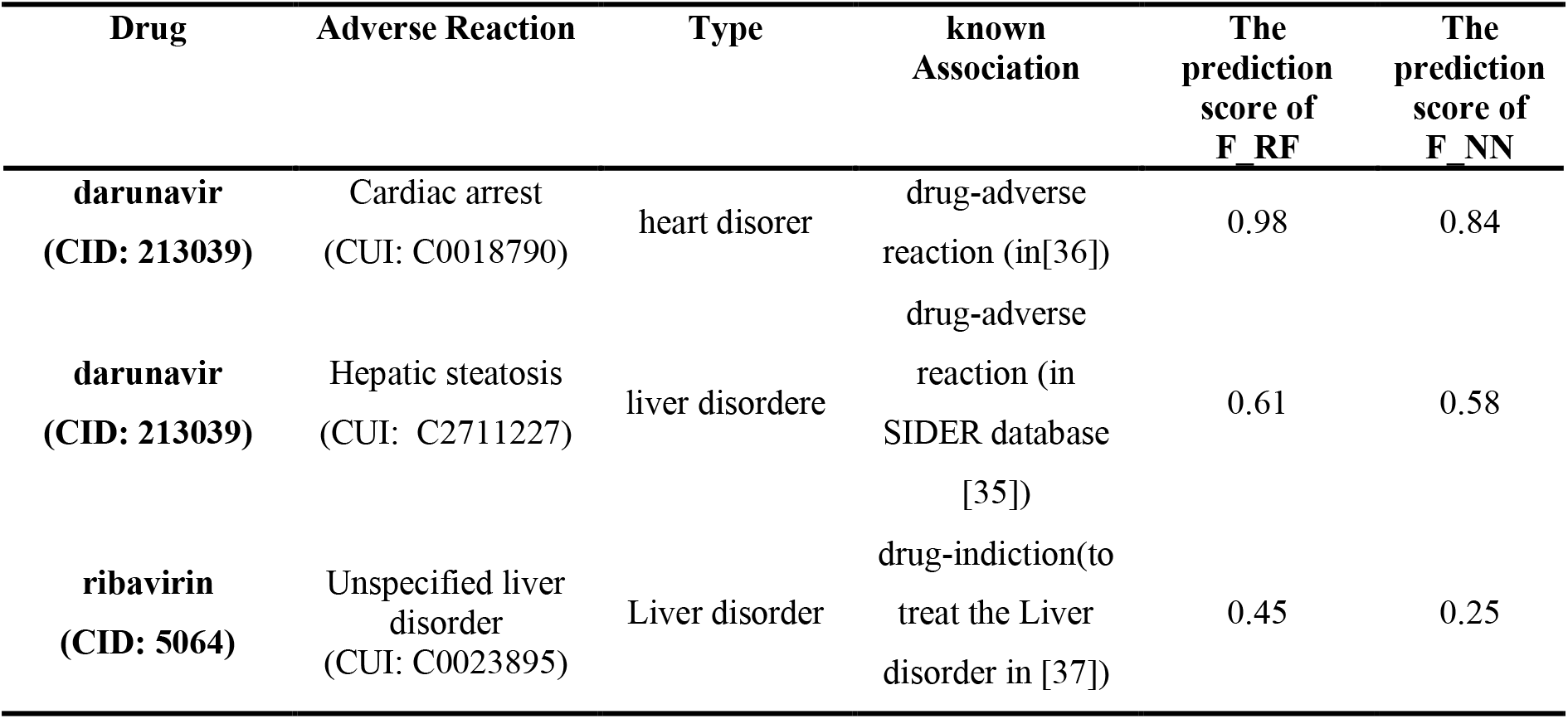
The results of three case studies.

| Drug | Adverse Reaction | Type | known Association | The prediction score of F_RF | The prediction score of F_NN |
| --- | --- | --- | --- | --- | --- |
| **darunavir**<br>(CID: 213039) | Cardiac arrest<br>(CUI: C0018790) | heart disorder | drug-adverse reaction (in[36]) | 0.98 | 0.84 |
| **darunavir**<br>(CID: 213039) | Hepatic steatosis<br>(CUI: C2711227) | liver disorder | drug-adverse reaction (in SIDER database [35]) | 0.61 | 0.58 |
| **ribavirin**<br>(CID: 5064) | Unspecified liver disorder<br>(CUI: C0023895) | Liver disorder | drug-indication(to treat the Liver disorder in [37]) | 0.45 | 0.25 |

Moreover, F_RF suggests drug ribavirin and adverse reaction Liver disorder as a negative association. It is mentioned in [37] which this drug can treat Liver disorder. Table 11 depicts the case studies and the results of F_NN and F_RF. F_NN confirms the results of F_RF.

## 5. Conclusion

This study proposed a framework called F_RF based on a random forest classifier to predict drug-adverse reaction associations. For this aim, a similarity vector is suggested by the drug-drug similarity score and the adverse reactions similarity function as the drug and adverse reaction representations. As the performance of machine learning methods depends on the training data, similarly to Zhang et al.[26], the drug-adverse reactions and drug-indication are considered positive and negative data, respectively. Then, another framework was introduced using a neural network named F_NN. Comparing the corresponding of these frameworks indicated the F_RF got higher evaluation scores than F_NN for predicting rare and non-rare adverse reactions. Later, two state-of-the-art matrix factorization methods, CS and TMF, were improved to *CS*^*Phen*^ and *TMF*_*Targ*_, *TMF*^*Phen*^ and 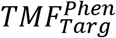 and contrast with F_RF. According to the results, F_RF is performed more accurately than these models. Moreover, F_RF framework was compared with some popular machine learning approaches in ADR problem. Although some methods exclude rare adverse reactions [24][19] or use more features to solve the ADR problem, F_RF utilized all drug-adverse reactions, including rare ones, and fewer features. The results announced that the F_RF performance was better than most of them. Meanwhile, F_RF correctly predicted cardiac arrest and hepatic steatosis as darunavir adverse reactions and suggested ribavirin to treat the liver disorder.

We conclude that using similarity vectors as drug and adverse reaction features and considering drug indications as negative data can improve drug-adverse reaction association prediction. Moreover, applying a random forest classifier with less computational complexity than other models achieves higher performance scores.

In the future, we aim to assess the 3D structures of drugs to increase the performance of drug-adverse reaction association prediction. In addition, applying drug-related clinical information can improve the accuracy of the model.

## Footnotes

1 https://www.mathworks.com/help/stats/treebagger.html

